# Bloodstream Infection with Extended-spectrum Beta-lactamase–producing *Escherichia coli*: the role of virulence genes

**DOI:** 10.1101/366187

**Authors:** Wan-Ting Hung, Ming-Fang Cheng, Fan-Chen Tseng, Yao-Shen Chen, Susan Shin-Jung Lee, Tsung-Hsien Chang, Hsi-Hsun Lin, Chih-Hsin Hung, Jiun-Ling Wang

## Abstract

**Background:** Various bacterial putative virulence factors are involved in the pathogenesis of bacterial infection. However, the effect of comorbidities or infection syndrome in the association of virulence factors and mortality remains inconclusive.

**Method:** This study addressed whether specific sequence type (ST) and virulence factors of extended-spectrum beta-lactamase–producing *Escherichia coli* (ESBL-EC) are associated with different outcomes in patients with bloodstream infection.121 adults from southern Taiwan with ESBL-producing *E. coli* bloodstream infections were enrolled during a 6-year period. Demographic data, including infection syndromes, underlying disease and outcomes, were collected. The virulence factors in isolates were analyzed by PCR and multilocus sequence typing.

**Result:** Positivity for the virulence genes i*ha*, *hlyD*, *sat*, *iut*, *fyu*, *malX*, *ompT*, *usp* and *traT* was associated with ST131 positivity (P<0.05). Some ESBL-EC virulence genes associated with urinary tract infection (UTI) were revealed. Positivity for ST405 and the virulence genes *iroN* and *iss* was significantly associated with increased 30-day mortality (death within 30 days) on univariate analysis (P<0.05). Independent risk factors of 30-day mortality in bacteremic patients with UTI included underlying chronic liver disease and malignancy. ST131 was borderline associated with 30-day mortality. Independent risk factors associated with 30-day mortality among bacteremic patients without UTI included comorbidities and *iroN* positivity.

**Conclusion:** In bacteremic patients with UTI, and the ST131 clone was borderline associated with mortality. Positivity for the virulence gene *iroN* may be linked to mortality in bacteremic patients without UTI.

## Introduction

Various proteins have been studied as putative virulence factors and shown to be involved in adhesion to and invasion of host cells, iron availability, toxic effects on host cells, or protection against the host’s immune system. Some virulence factors are associated with urinary tract infection (UTI) and some are associated with blood invasion from the bowel. Previous study revealed *pap*, *malX*, *usp*, *fyuA* and phylogentic group B genes positively associated with mortality.[1] The co-occurrence of multiple genes encoding *papC*, *sfa*, *usp* and *cnf1* virulence factors probably predisposes *Escherichia coli* to translocation from the gastrointestinal tract to the vascular bed in patients with hematologic malignancies.[2] A recent study showed that differences in virulence gene prevalence between cystitis and pyelonephritis isolates could be limited to 8 genes.[3] However, the association of virulence gene and disease severity is still controversial.[4, 5] Few studies have focused on the relationship of virulence factors and mortality in extended-spectrum beta-lactamase (ESBL) *E. coli* (ESBL-EC) bacteremia.[6, 7] As well, these studies usually lack information on sequence type (ST) or comorbidities.

From 2000 to 2006, *E. coli* clone ST131 producing CTX-M-15 ESBL has been identified in 3 continents.[8, 9] In most Asian countries outside India including Taiwan, the CTX-M-14 type was the most prevalent ESBL among *E. coli* isolates. In addition, ST131-producing CTX-M-15 has been found prevalent in Japan, with rare cases described in Indonesia and China.[10-13] Until this report, a detailed report of virulence data related to ST131 in Taiwan was lacking. In our previous study, we found ST131 in about one third of patients with bacteremia and ESBL-EC.[10] Our study cohort showed ST131 isolates in both community- and hospital-onset infections. The virulence gene distribution in CTX-M-14 ESBL-EC with ST131 is unknown. In addition, we do not know whether other STs (e.g., ST38, ST405, ST95, or ST69) are associated with mortality in ESBL-EC bacteremia.

This study aimed to understand the association between specific STs and virulence factors in patients with ESBL-EC bloodstream infection, in particular, any association with comorbidities and infection syndrome.

## Methods

### Patients and ethics statement

The study consisted of adults aged 16 years or older with ESBL-EC bloodstream infection in a teaching hospital in southern Taiwan during 2005-2010.

All patients were evaluated by using a structured recording form. Each clinical course of infection was evaluated and recorded according to the information supplied by primary care physicians and medical records. The diagnosis of the infection focus of bacteremia was based on clinical, bacteriological, and radiological investigations. If no infection focus could be identified, the bacteremia was classified as primary infection. Only strains from the first bacteremic episode were included in the analysis.

Demographic data, including infection syndrome, underlying disease and outcome, were collected from medical charts. The following items were recorded for each patient: age; sex; underlying illness; severity of illness (classified by Charlson comorbidity score); history of hospitalization or outpatient department involvement; antibiotic use history for more than 7 days before the bacteremic episode; surgical history within the previous 3 months; existence of a nasogastric tube, central venous catheter, or urinary catheter; initial empirical antimicrobial agents used; and outcome. If an *in vitro* active antimicrobial agent was administered before the final blood culture result, it was considered adequate empirical therapy. This study was approved by the institutional review board (IRB) at E-Da Hospital (no. EMRP-098-006).

Written informed consents were waived by IRB due to no harms or minimal risk in the patients in this study.

### Virulent gene detection

The virulence factors for ESBL-EC were analyzed by PCR and multilocus sequence typing (MLST). Virulent genes examined included *iutA* (aerobactin iron transport system), *kpsMTII* (capsular polysaccharide), *hlyA* (hemolysin), *sfa/foc* (S fimbriae and F1C fimbriae), *afa/draBC* (Dr family adhesin) and *Pap* operon (pyelonephritis-associated Pili). PCR amplification involved a 25-μL reaction mixture containing template DNA (2 μL boiled lysate, 4 mM MgCl_2_, 0.8 mM each of 4 dNTPs, 0.6 μM each primer [concentration 0.3 μM], and 2.5 units AmpliTaq Gold in 1× PCR buffer [Perkin Elmer, Branchburg, NJ]). The primer use was as described.[14] MLST was conducted according to the website http://mlst.warwick.ac.uk/mlst/dbs/Ecoli.

### Statistical analyses

Analyses involved use of SAS 9.3 (Cary, NC, USA) and all statistical tests were two-tailed. Continuous variables are described and were compared by Mann-Whitney U test. Categorical variables are described with percentages and were compared by Fisher exact test. Factors that may have a dose–response or trend relationship were tested by Cochran–Armitage test for trend. Previous studies suggested that the bacterial strain and site of infections may have important effects on the severity of diseases, so the following characteristics were compared first in the analyses: (1) *E. coli* O25b-ST131 versus non-O25b-ST131, (2) UTI versus non-UTI, and (3) death within 30 days versus survival > 30 days. Possible interactions or effect modification for these 3 characteristics with other patients or bacterial factors were also examined by stratified analyses with the Breslow-Day heterogeneity test, with significance set at p<0.1. If a significant difference was found, subsequent analyses would be stratified by this factor. Factors showing significance at p<0.1 on univariate analysis were all considered in the multivariate analysis. Stratified logistic regression was used to estimate independent odds ratios (ORs) and 95% confidence intervals (95% CIs) for bacterial strain, virulence factor or patient characteristics. P<0.05 was considered statistically significant.

## Results

### Virulence gene distribution in ESBL-EC and between UTI and non-UTI bacteremic patients

For the 121 adults included in the study, we analyzed the virulence genes of adhesions, toxins, sideropherores, capsule and miscellaneous distribution with the ST131 and non-ST131 clone. The mortality of different ST type was shown in Figuire 1. The comorbidity of ST131 vs. ST131 was shown in Supplement Table 1. Positivity for the genes *iha*, *hlyD*, *sat*, *iutA*, *fyuA*, *malX*, *ompT*, and *traT* was more frequent with the ST131 than non-ST131 clone (P<0.05) and that for *vat*, *kpsMTII*, *kpsMTIIK1* and *traT* was more frequent with the non-ST131 clone (P<0.05) (Table 1). Positivity for the virulence genes *pap*, *iha*, *hlyD*, *sat*, *iutA*, *fyuA*, *malX*, *ompT*, *usp*, and *traT* was more frequent with than without UTI (Table 2).

**Table 1:** Virulence gene distribution for patients with ST131 and non-ST131 clone

| Virulence gene | Total<br>n=121 | ST131<br>n=36 | Non-ST131<br>n=85 | *P* | OR (95% CI) |
| --- | --- | --- | --- | --- | --- |
| Adhesins |  |  |  |  |  |
| *papA* | 44(36.4) | 13(36.1) | 31(36.5) | 0.97 | 1.0(0.43~2.21) |
| *fimH* | 117(96.7) | 35(97.2) | 82(96.5) | 0.83 | 1.3(0.12~12.7) |
| *papEF* | 43(35.5) | 13(36.1) | 30(35.3) | 0.93 | 1.0(0.46~2.33) |
| *papG* allele II | 35(28.9) | 14(38.9) | 21(24.7) | 0.12 | 1.9(0.84~4.45) |
| *papG* allele III | 9(7.4) | 1(2.8) | 8(9.4) | 0.20 | 3.6(0.43~30.20) |
| *tsh* | 6(5.0) | 1(2.8) | 5(5.9) | 0.47 | 0.5(0.05~4.05) |
| *iha* | 61(50.4) | 31(86.1) | 30(35.3) | **<0.01** | 11.4(4.00~32.29) |
| Toxins |  |  |  |  |  |
| *hlyD* | 25(20.7) | 12(33.3) | 13(15.3) | **0.03** | 2.8(1.11~6.88) |
| *sat* | 55(45.5) | 27(75.0) | 28(32.9) | **<0.01** | 6.1(2.53~14.71) |
| *vat* | 16(13.2) | 1(2.8) | 15(17.6) | **0.02** | 0.1(0.01~1.05) |
| Siderophores |  |  |  |  |  |
| *iroN* | 23(19.0) | 3(8.3) | 20(23.5) | 0.05 | 0.3(0.08~1.06) |
| *iutA* | 97(80.2) | 34(94.4) | 63(74.1) | **0.01** | 5.9(1.31~26.77) |
| *fyuA* | 96(79.3) | 34(94.4) | 62(72.9) | **0.01** | 6.3(1.40~28.38) |
| Capsule |  |  |  |  |  |
| *kpsMTII* | 70(57.9) | 15(41.7) | 55(64.7) | **0.02** | 0.4(0.17~0.86) |
| *kpsMTIIK1* | 22(18.2) | 1(2.8) | 21(24.7) | **<0.01** | 0.9(0.11~0.67) |
| *kpsMTIIK5* | 43(35.5) | 12(33.3) | 31(36.5) | 0.74 | 0.9(0.38~1.98) |
| Miscellaneous |  |  |  |  |  |
| *malX* | 69(57.0) | 31(86.1) | 38(44.7) | **<0.01** | 7.7(2.71~21.62) |
| *ompT* | 67(55.4) | 32(88.9) | 35(41.2) | **<0.01** | 11.4(3.70~35.22) |
| *iss* | 14(11.6) | 3(8.3) | 11(12.9) | 0.47 | 0.6(0.16~2.33) |
| *traT* | 101(83.5) | 34(94.4) | 67(78.8) | **0.03** | 4.6(1.00~20.84) |

Data are no. (%).

**Table 2:** Virulence gene distribution for UTI and non-UTI bacteremic patients

| Type of infection | Total<br>n=121 | UTI<br>n=65 | Non-UTI<br>n=56 | *P* | OR (95% CI) |
| --- | --- | --- | --- | --- | --- |
| Adhesins |  |  |  |  |  |
| *papA* | 44(36.4%) | 33(50.8%) | 11(19.6%) | <0.01 | 4.21(1.86-9.57) |
| *papEF* | 43(35.5%) | 33(50.8%) | 10(17.9%) | <0.01 | 4.74(2.50-10.97) |
| *papG* allele II | 35(28.9%) | 29(44.6%) | 6(10.7%) | <0.01 | 6.71(2.53-17.84) |
| *iha* | 61(50.4%) | 44(67.7%) | 17(30.4%) | <0.01 | 4.80(2.22-10.39) |
| Toxins |  |  |  |  |  |
| *hlyD* | 25(20.7%) | 20(30.8%) | 5(8.9%) | <0.01 | 4.53(1.57-13.06) |
| *sat* | 55(45.5%) | 40(61.5%) | 15(26.8%) | <0.01 | 4.37(2.01-9.48) |
| Siderophores |  |  |  |  |  |
| *iutA* | 97(80.2%) | 60(92.3%) | 37(66.1%) | <0.01 | 6.16(2.12-17.91) |
| *fyuA* | 96(79.3%) | 38(89.2%) | 38(67.9%) | 0.004 | 3.92(1.49-10.29) |
| Miscellaneous |  |  |  |  |  |
| *malX* | 69(57.0%) | 45(69.2%) | 24(42.9%) | 0.003 | 3.00(1.42-6.33) |
| *ompT* | 67(55.4%) | 46(70.8%) | 21(37.5%) | <0.01 | 4.03(1.88-8.63) |
| *usp* | 56(46.3%) | 36(55.4%) | 20(35.7%) | 0.03 | 2.23(1.07-4.65) |
| *traT* | 101(83.5%) | 60(92.3%) | 41(73.2%) | 0.005 | 4.39(1.48-13.02) |

Data are no. (%).
UTI: urinary tract infection

### Virulence gene survey in patients with 30-day mortality and survival > 30 days

Mortality within 30 days was associated more with ST131 than ST405 infection (28% vs. 15.6%). Positivity for most virulence genes was not associated with 30-day mortality, but 2 virulence genes, siderophores *iroN* and *iss*, were highly associated with 30-day mortality on univariate analysis (OR 3.36 [95% CI 1.30-8.68] and 3.23 [1.05-10.24], P<0.05) (Table 3).

**Table 3:**
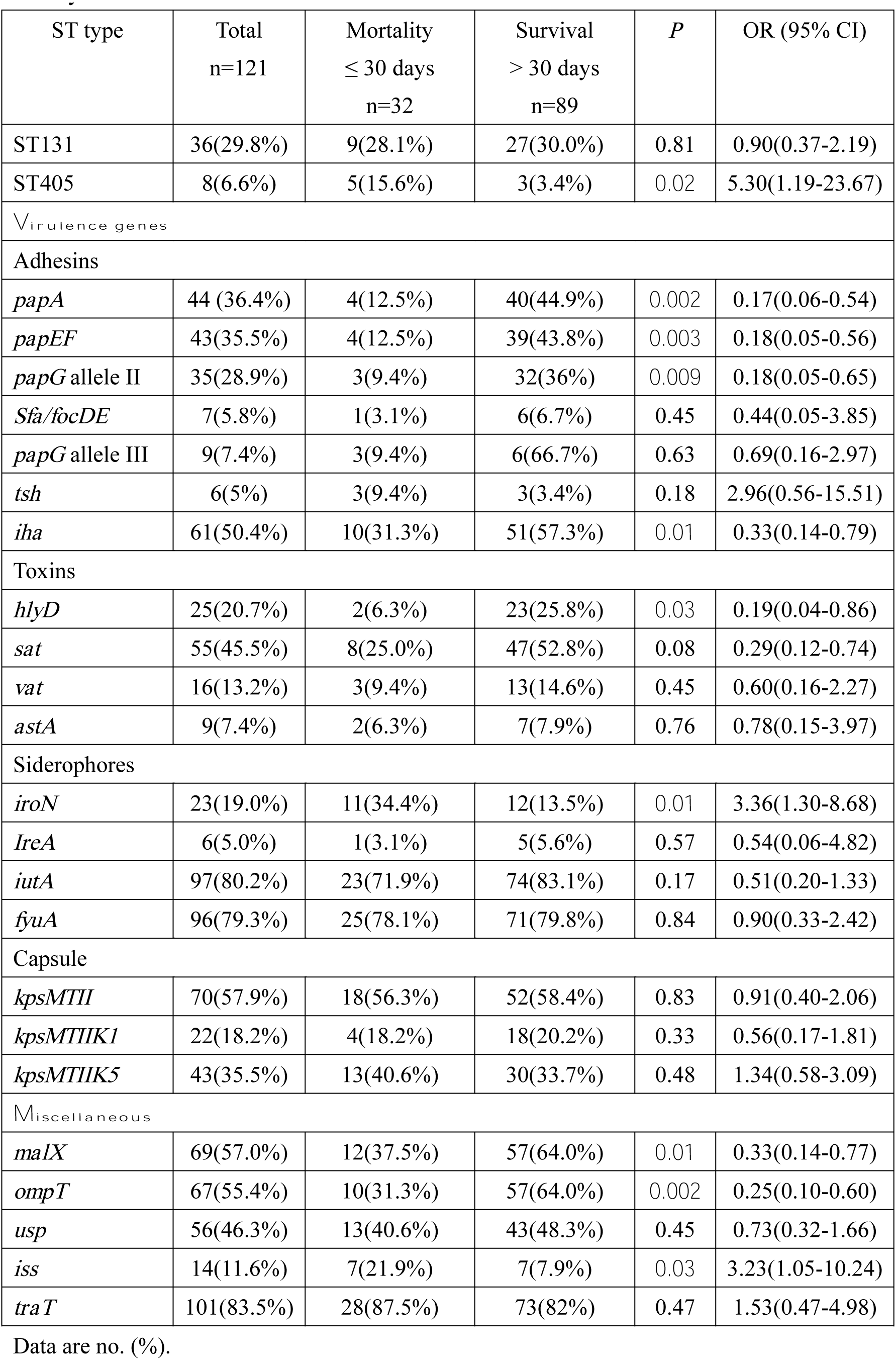
Virulence gene distribution for patients with 30-day mortality and survival > 30 days

| ST type | Total<br>n=121 | Mortality<br>≤ 30 days<br>n=32 | Survival<br>> 30 days<br>n=89 | *P* | OR (95% CI) |
| --- | --- | --- | --- | --- | --- |
| ST131 | 36(29.8%) | 9(28.1%) | 27(30.0%) | 0.81 | 0.90(0.37-2.19) |
| ST405 | 8(6.6%) | 5(15.6%) | 3(3.4%) | **0.02** | 5.30(1.19-23.67) |
| **Virulence genes** |  |  |  |  |  |
| Adhesins |  |  |  |  |  |
| *papA* | 44 (36.4%) | 4(12.5%) | 40(44.9%) | **0.002** | 0.17(0.06-0.54) |
| *papEF* | 43(35.5%) | 4(12.5%) | 39(43.8%) | **0.003** | 0.18(0.05-0.56) |
| *papG* allele II | 35(28.9%) | 3(9.4%) | 32(36%) | **0.009** | 0.18(0.05-0.65) |
| *Sfa/focDE* | 7(5.8%) | 1(3.1%) | 6(6.7%) | 0.45 | 0.44(0.05-3.85) |
| *papG* allele III | 9(7.4%) | 3(9.4%) | 6(6.7%) | 0.63 | 0.69(0.16-2.97) |
| *tsh* | 6(5%) | 3(9.4%) | 3(3.4%) | 0.18 | 2.96(0.56-15.51) |
| *iha* | 61(50.4%) | 10(31.3%) | 51(57.3%) | **0.01** | 0.33(0.14-0.79) |
| Toxins |  |  |  |  |  |
| *hlyD* | 25(20.7%) | 2(6.3%) | 23(25.8%) | **0.03** | 0.19(0.04-0.86) |
| *sat* | 55(45.5%) | 8(25.0%) | 47(52.8%) | 0.08 | 0.29(0.12-0.74) |
| *vat* | 16(13.2%) | 3(9.4%) | 13(14.6%) | 0.45 | 0.60(0.16-2.27) |
| *astA* | 9(7.4%) | 2(6.3%) | 7(7.9%) | 0.76 | 0.78(0.15-3.97) |
| Siderophores |  |  |  |  |  |
| *iroN* | 23(19.0%) | 11(34.4%) | 12(13.5%) | **0.01** | 3.36(1.30-8.68) |
| *IreA* | 6(5.0%) | 1(3.1%) | 5(5.6%) | 0.57 | 0.54(0.06-4.82) |
| *iutA* | 97(80.2%) | 23(71.9%) | 74(83.1%) | 0.17 | 0.51(0.20-1.33) |
| *fyuA* | 96(79.3%) | 25(78.1%) | 71(79.8%) | 0.84 | 0.90(0.33-2.42) |
| Capsule |  |  |  |  |  |
| *kpsMTII* | 70(57.9%) | 18(56.3%) | 52(58.4%) | 0.83 | 0.91(0.40-2.06) |
| *kpsMTIIK1* | 22(18.2%) | 4(18.2%) | 18(20.2%) | 0.33 | 0.56(0.17-1.81) |
| *kpsMTIIK5* | 43(35.5%) | 13(40.6%) | 30(33.7%) | 0.48 | 1.34(0.58-3.09) |
| **Miscellaneous** |  |  |  |  |  |
| *malX* | 69(57.0%) | 12(37.5%) | 57(64.0%) | **0.01** | 0.33(0.14-0.77) |
| *ompT* | 67(55.4%) | 10(31.3%) | 57(64.0%) | **0.002** | 0.25(0.10-0.60) |
| *usp* | 56(46.3%) | 13(40.6%) | 43(48.3%) | 0.45 | 0.73(0.32-1.66) |
| *iss* | 14(11.6%) | 7(21.9%) | 7(7.9%) | **0.03** | 3.23(1.05-10.24) |
| *traT* | 101(83.5%) | 28(87.5%) | 73(82%) | 0.47 | 1.53(0.47-4.98) |

Data are no. (%).

### Factors associated with 30-day mortality with or without UTI

Independent risk factors associated with 30-day mortality among 65 bacteremia patients with UTI included underlying chronic liver disease and malignancy (OR 7.38 [95% CI 1.5-36.3], p<0.05). ST131 was borderline associated with 30-day mortality in bacteremia patients with UTI (OR 10.8 [1-117.7], P=0.05). Positivity for *malX* was a protective factor (OR 0.033 [0.03-0.357]). Independent risk factors associated with 30-day mortality among 56 bacteremic patients without UTI included underlying chronic liver disease and malignancy (OR 8.66 [1.7-42.63]) and positivity for *iroN* (OR 5.88 [1.39-24.9], p<0.05) (Table 4).

**Table 4:**
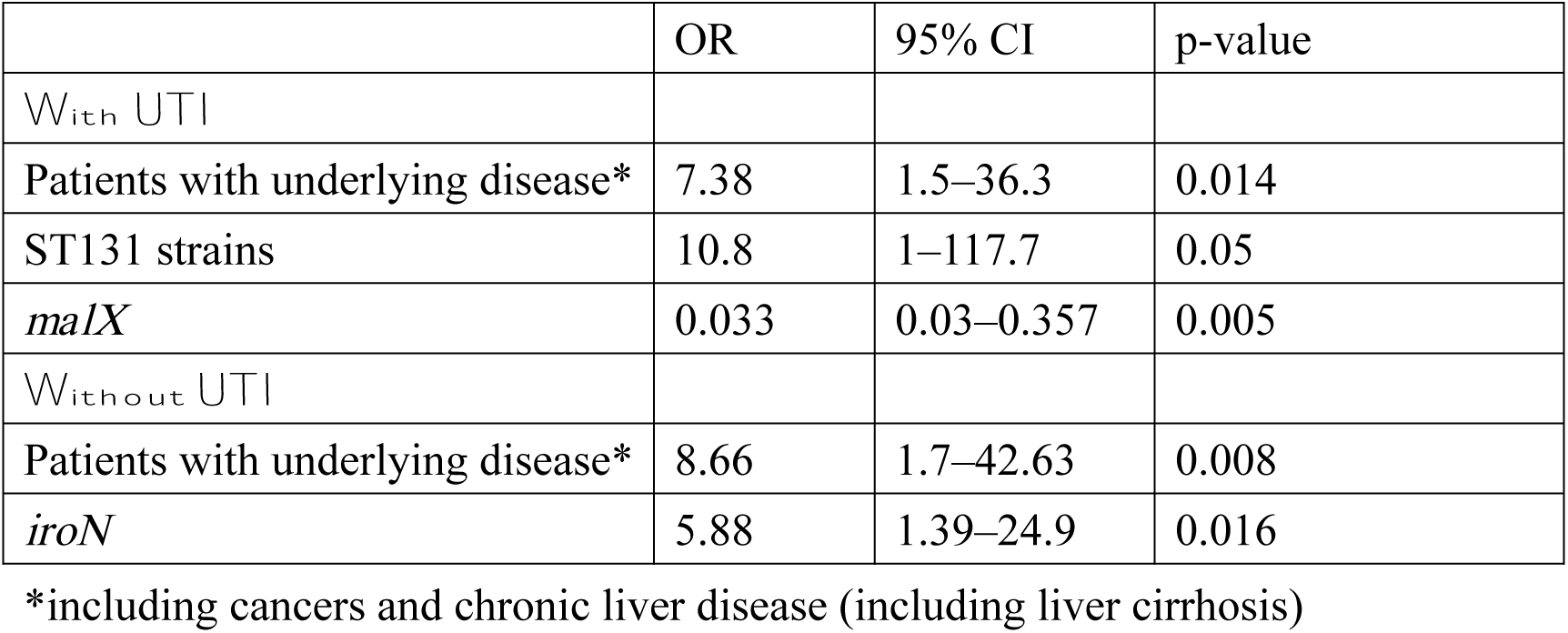
Multivariate regression analysis of factors associated with 30-day mortality in patients with UTI (n=65) and without UTI (n=56)

## Discussion

We performed a mortality analysis in a cohort of ESBL-EC bacteremic patients and considered host factors such as underlying disease and infection type, bacteria characteristics including ST, and virulence genes. Positivity for the siderophores *iroN* and miscellaneous *iss* genes was associated with 30-day mortality in patients with ESBL-EC bacteremia. In particular, positivity for the iron gene was associated with 30-day mortality in non-UTI bacteremic patients in stratified logistic regression analysis.

The virulence score is greater with the ST131 clone than non-ST131 clone in isolates.[10] Our virulence gene analysis indicated that clone ST131 strains were more like to be associated with *traT*, *iutA*, *iha*, *ompT*, *fimH*, *malX*, *fyuA*, *kpsM*, and *sat* genes than non-ST131 strains.

Particularly, the virulence gene response for hemolysin and siderophores was predominant in ST131 strains. This finding is consistent with its high virulence potential in a mouse model of septicemia[15] and perhaps explains in part the higher epidemiological success of the ST131 clonal group.

The prevalence of P fimbriation (*papA*) in isolates from patients with bacteremia with UTI (urosepsis and pyelonephritis) is high as in isolates from patients with bacteremia from other sources. P fimbriae may contribute to the ability of *E. coli* strains to cause UTI.[16]

The aerobactin system (*iutA*), responses for siderophores in *E. coli*, is encoded by a five-gene operon, which is commonly found together with P fimbriae in isolates from patients with UTI and urosepsis. An association of chromosomal aerobactin with hemolysin (hlyD, and *hlyA*) is found among urosepsis isolates. Hemolytic uropathogenic strains almost always also express P fimbriae (*papA*). Hemolysin production is associated with the human pathogenic strain of *E. coli*, especially those causing more clinically severe forms of UTI.[16] This finding is similar to our observation of the virulence gene response for adhesion genes (*papA*, *papEF* and *papG allele II*), toxins such as hemolysin (*hlyD*), and siderophores (*iutA*, and *fyuA*) associated with a specific infection syndrome such as UTI (Table 2).

A French cohort study revealed an association of positivity for *papGIII*, septic shock at baseline and a non-urinary tract origin of sepsis with a fatal outcome.[17] A cohort of cirrhotic patients with *E. coli* bacteremia and spontaneous bacterial peritonitis showed no significant association between specific clonal groups and patient characteristics, type of infection, or outcome.[18] Mora-Rillo et al. found positivity for *fyuA* associated with increased mortality, and that of P fimbriae genes had a protective role.[19] A study in Spain of ESBL *E coli* bacteremia showed *ibeA* associated with increased mortality, with *papG alleleII* having a protective effect.[6]

We found *iroN* and *iss* more frequently associated with 30-day mortality than other virulence genes on univariate analysis. However, we also found positivity for *papA*, *papEF*, *papG alleleII*, *iha*, *hlyD*, *sat* and *malX* with a protective effect on mortality. *FyuA* and *papGIII* genes had no significant impact on 30-day mortality.

The virulence gene *iroN* is a novel *E. coli* gene, with 77% DNA homology to a catecholate siderophore receptor gene recently identified in Salmonella. It is linked to the P-pilus (*prs*) and F1C fimbrial (*foc*) gene clusters on a pathogenicity island and appears to have been acquired by IS1230-mediated horizontal transmission.[19] Johnson suggested that *iroN* of *E. coli* could be a good target for an anti-virulence factor (VF) intervention among immunocompromised and non-immunocompromised hosts because the prevalence of *iroN* of *E. coli* was increased in the presence of host compromise, and *iroN* of *E. coli* could also be an effective target for anti-VF intervention among multidrug-resistant *E. coli*.[20] Russo et al., in a mouse model of ascending UTI, showed that *iroN* positivity contributed significantly to the ability to colonize the mouse bladder, kidneys, and urine, evidence that *iroN* is a urovirulence factor.[21] Dozois et al. performed specific deletion of the aerobactin siderophore system and *E. coli iroN* locus and demonstrated that these pathogen-specific systems contribute to the virulence of avian strains.[22] Nègre et al. found that the *iroN* gene itself played a key role in the virulence of *E. coli* strain C5, possibly by contributing to the sustained high-level bacteremia that precedes meningitis.[23]

Our data showed *iroN* indeed associated with higher 30-day mortality on univariate analysis and in patients with cancers and chronic liver disease including liver cirrhosis, was associated with 30-day mortality without UTI.

There were several limitations in this study. First, the mortality analysis may be biased by selected isolates in single center in Taiwan. As well, the different strains may not be generalized to other parts of the world. Second, although we considered host factors in out multivariate analysis, we cannot exclude the possibility that some virulence genes were linked to some specific hosts.

In conclusion, in adult ESBL *E.coli* bloodstream infection, some specific virulent genes were linked to specific ST and infection syndromes. Besides the host factors, the presence of virulence gene *iroN* may be linked to increased mortality in bacteremic patients without UTI.

## Declarations

## Acknowledgements

nil

## Abbreviation

(ST): specific sequence type
(ESBL-EC): extended-spectrum beta-lactamase–producing Escherichia coli
(UTI): urinary tract infection
(MLST): multilocus sequence typing

---

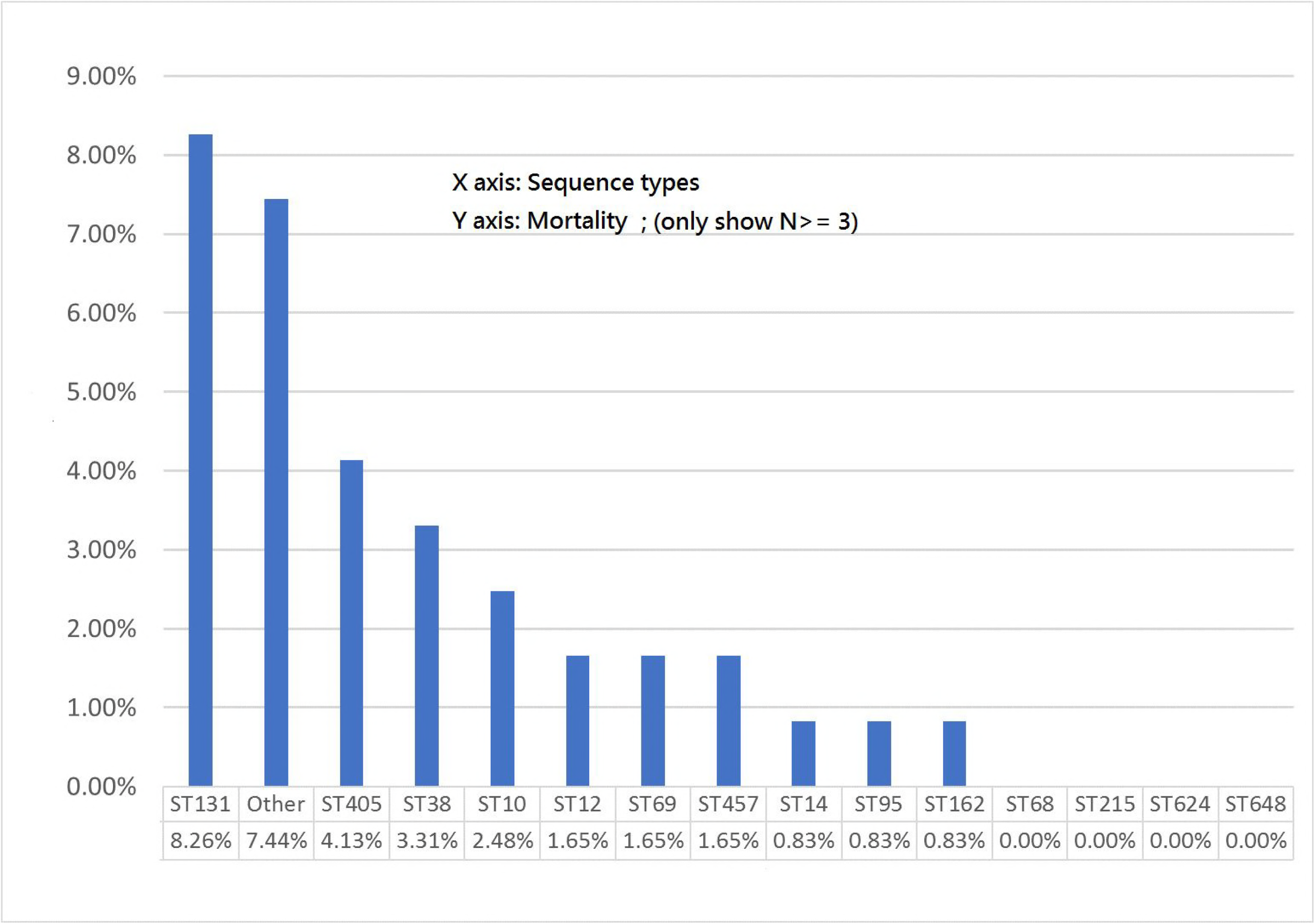

## References

1. Johnson JR, Clermont O, Menard M, Kuskowski MA, Picard B, Denamur E: Experimental mouse lethality of Escherichia coli isolates, in relation to accessory traits, phylogenetic group, and ecological source. The Journal of infectious diseases 2006, 194(8):1141-1150.

2. Krawczyk B, Sledzinska A, Szemiako K, Samet A, Nowicki B, Kur J: Characterisation of Escherichia coli isolates from the blood of haematological adult patients with bacteraemia: translocation from gut to blood requires the cooperation of multiple virulence factors. European journal of clinical microbiology & infectious diseases: official publication of the European Society of Clinical Microbiology 2015, 34(6):1135-1143.

3. Kudinha T, Johnson JR, Andrew SD, Kong F, Anderson P, Gilbert GL: Distribution of phylogenetic groups, sequence type ST131, and virulence-associated traits among Escherichia coli isolates from men with pyelonephritis or cystitis and healthy controls. Clinical microbiology and infection: the official publication of the European Society of Clinical Microbiology and Infectious Diseases 2013, 19(4):E173-180.

4. Landraud L, Jaureguy F, Frapy E, Guigon G, Gouriou S, Carbonnelle E, Clermont O, Denamur E, Picard B, Lemichez E et al: Severity of Escherichia coli bacteraemia is independent of the intrinsic virulence of the strains assessed in a mouse model. Clinical microbiology and infection: the official publication of the European Society of Clinical Microbiology and Infectious Diseases 2013, 19(1):85-90.

5. Picard B, Garcia JS, Gouriou S, Duriez P, Brahimi N, Bingen E, Elion J, Denamur E: The link between phylogeny and virulence in Escherichia coli extraintestinal infection. Infection and immunity 1999, 67(2):546-553.

6. Rodriguez-Bano J, Mingorance J, Fernandez-Romero N, Serrano L, Lopez-Cerero L, Pascual A: Outcome of bacteraemia due to extended-spectrum beta-lactamase-producing Escherichia coli: impact of microbiological determinants. The Journal of infection 2013, 67(1):27-34.

7. Rodriguez-Bano J, Mingorance J, Fernandez-Romero N, Serrano L, Lopez-Cerero L, Pascual A: Virulence profiles of bacteremic extended-spectrum beta-lactamase-producing Escherichia coli: association with epidemiological and clinical features. PloS one 2012, 7(9):e44238.

8. Coque TM, Novais A, Carattoli A, Poirel L, Pitout J, Peixe L, Baquero F, Canton R, Nordmann P: Dissemination of clonally related Escherichia coli strains expressing extended-spectrum beta-lactamase CTX-M-15. Emerg Infect Dis 2008, 14(2):195-200.

9. Peirano G, Pitout JD: Molecular epidemiology of Escherichia coli producing CTX-M beta-lactamases: the worldwide emergence of clone ST131 O25:H4. International journal of antimicrobial agents 2010, 35(4):316-321.

10. Chung HC, Lai CH, Lin JN, Huang CK, Liang SH, Chen WF, Shih YC, Lin HH, Wang JL: Bacteremia caused by extended-spectrum-beta-lactamase-producing Escherichia coli sequence type ST131 and non-ST131 clones: comparison of demographic data, clinical features, and mortality. Antimicrob Agents Chemother 2012, 56(2):618-622.

11. Kim SY, Park YJ, Johnson JR, Yu JK, Kim YK, Kim YS: Prevalence and characteristics of Escherichia coli sequence type 131 and its H30 and H30Rx subclones: a multicenter study from Korea. Diagnostic microbiology and infectious disease 2016, 84(2):97-101.

12. Hussain A, Ranjan A, Nandanwar N, Babbar A, Jadhav S, Ahmed N: Genotypic and phenotypic profiles of Escherichia coli isolates belonging to clinical sequence type 131 (ST131), clinical non-ST131, and fecal non-ST131 lineages from India. Antimicrobial agents and chemotherapy 2014, 58(12):7240-7249.

13. Park HJ, Lee YM, Bang KM, Park SY, Moon SM, Park KH, Chong YP, Kim SH, Lee SO, Choi SH et al: Clinical significance of Staphylococcus aureus bacteremia in patients with liver cirrhosis. Eur J Clin Microbiol Infect Dis 2012, 31(12):3309-3316.

14. Johnson JR, Stell AL: Extended virulence genotypes of Escherichia coli strains from patients with urosepsis in relation to phylogeny and host compromise. The Journal of infectious diseases 2000, 181(1):261-272.

15. Clermont O, Lavollay M, Vimont S, Deschamps C, Forestier C, Branger C, Denamur E, Arlet G: The CTX-M-15-producing Escherichia coli diffusing clone belongs to a highly virulent B2 phylogenetic subgroup. The Journal of antimicrobial chemotherapy 2008, 61(5):1024-1028.

16. Johnson JR: Virulence factors in Escherichia coli urinary tract infection. Clinical microbiology reviews 1991, 4(1):80-128.

17. Jaureguy F, Carbonnelle E, Bonacorsi S, Clec’h C, Casassus P, Bingen E, Picard B, Nassif X, Lortholary O: Host and bacterial determinants of initial severity and outcome of Escherichia coli sepsis. Clinical microbiology and infection: the official publication of the European Society of Clinical Microbiology and Infectious Diseases 2007, 13(9):854-862.

18. Bert F, Johnson JR, Ouattara B, Leflon-Guibout V, Johnston B, Marcon E, Valla D, Moreau R, Nicolas-Chanoine MH: Genetic diversity and virulence profiles of Escherichia coli isolates causing spontaneous bacterial peritonitis and bacteremia in patients with cirrhosis. Journal of clinical microbiology 2010, 48(8):2709-2714.

19. Mora-Rillo M, Fernandez-Romero N, Francisco CN, Diez-Sebastian J, Romero-Gomez MP, Fernandez FA, Lopez JR, Mingorance J: Impact of virulence genes on sepsis severity and survival in Escherichia coli bacteremia. Virulence 2015, 6(1):93-100.

20. Johnson JR, Russo TA, Tarr PI, Carlino U, Bilge SS, Vary JC, Jr., Stell AL: Molecular epidemiological and phylogenetic associations of two novel putative virulence genes, iha and iroN(E. coli), among Escherichia coli isolates from patients with urosepsis. Infection and immunity 2000, 68(5):3040-3047.

21. Russo TA, McFadden CD, Carlino-MacDonald UB, Beanan JM, Barnard TJ, Johnson JR: IroN functions as a siderophore receptor and is a urovirulence factor in an extraintestinal pathogenic isolate of Escherichia coli. Infection and immunity 2002, 70(12):7156-7160.

22. Dozois CM, Daigle F, Curtiss R, 3rd: Identification of pathogen-specific and conserved genes expressed in vivo by an avian pathogenic Escherichia coli strain. Proceedings of the National Academy of Sciences of the United States of America 2003, 100(1):247-252.

23. Negre VL, Bonacorsi S, Schubert S, Bidet P, Nassif X, Bingen E: The siderophore receptor IroN, but not the high-pathogenicity island or the hemin receptor ChuA, contributes to the bacteremic step of Escherichia coli neonatal meningitis. Infection and immunity 2004, 72(2):1216-1220.

